# RNA regulates Glycolysis and Embryonic Stem Cell Differentiation via Enolase 1

**DOI:** 10.1101/2020.10.14.337444

**Authors:** Ina Huppertz, Joel I. Perez-Perri, Panagiotis Mantas, Thileepan Sekaran, Thomas Schwarzl, Lyudmila Dimitrova-Paternoga, Janosch Hennig, Pierre A. Neveu, Matthias W. Hentze

## Abstract

Cells must coordinate their metabolism and fate trajectories (*1, 2*), but the underlying mechanisms are only beginning to be discovered. To understand why the glycolytic enzyme enolase 1 (ENO1) binds RNA (*3–6*), we studied this phenomenon in vitro, in human cells, and during mouse embryonic stem cell differentiation. We find specific cellular RNA ligands that inhibit ENO1’s enzymatic activity in vitro. Increasing the concentration of these ligands in cultured cells inhibits glycolysis. We demonstrate that pluripotent stem cells expressing an ENO1 mutant that is hyper-inhibited by RNA are severely impaired in their glycolytic capacity and in endodermal differentiation, whereas cells with an RNA binding-deficient ENO1 mutant display disproportionately high endodermal marker expression. Our findings uncover ENO1 riboregulation as a novel form of metabolic control. They also describe an unprecedented mechanism involved in the regulation of stem cell differentiation.

**One Sentence Summary:** RNA directly regulates enzyme activity to control metabolism and stem cell fate

---

Glycolysis represents a classic metabolic pathway that has received renewed interest in recent years due to its role in stem cell differentiation and cancer biology (*1, 2*). Our fundamental understanding of glycolysis as a protein- and metabolite-controlled process has remained widely unchanged over the last decades. Intriguingly, the glycolytic enzyme enolase has been reproducibly found amongst the pool of proteins that associate with RNA in cultured mammalian cells and other organisms (*3–6*), but apart from enolase’s involvement in bacterial RNA degradation (*7, 8*) and its role in tRNA targeting to the mitochondria in yeast (*9, 10*), relatively limited functional insights have been gained, especially in mammalian systems.

Enolase 1 (ENO1) catalyzes the reversible interconversion between 2-phosphoglycerate (2-PG) and phosphoenolpyruvate (PEP). ENO1 has previously been shown to bind RNA in different organisms including rat (*11*), bacteria (*7, 8*) and yeast (*9, 10*). We first confirmed, by a PNK assay (*12*), that human ENO1 binds RNA in HeLa cells (Fig. 1A). In addition, we could estimate that under basal conditions around 10% of HeLa cell ENO1 is sensitive to RNase treatment when exposing RNase-treated or untreated lysates to sucrose density gradient centrifugation (Fig. 1B).

**Fig. 1.**
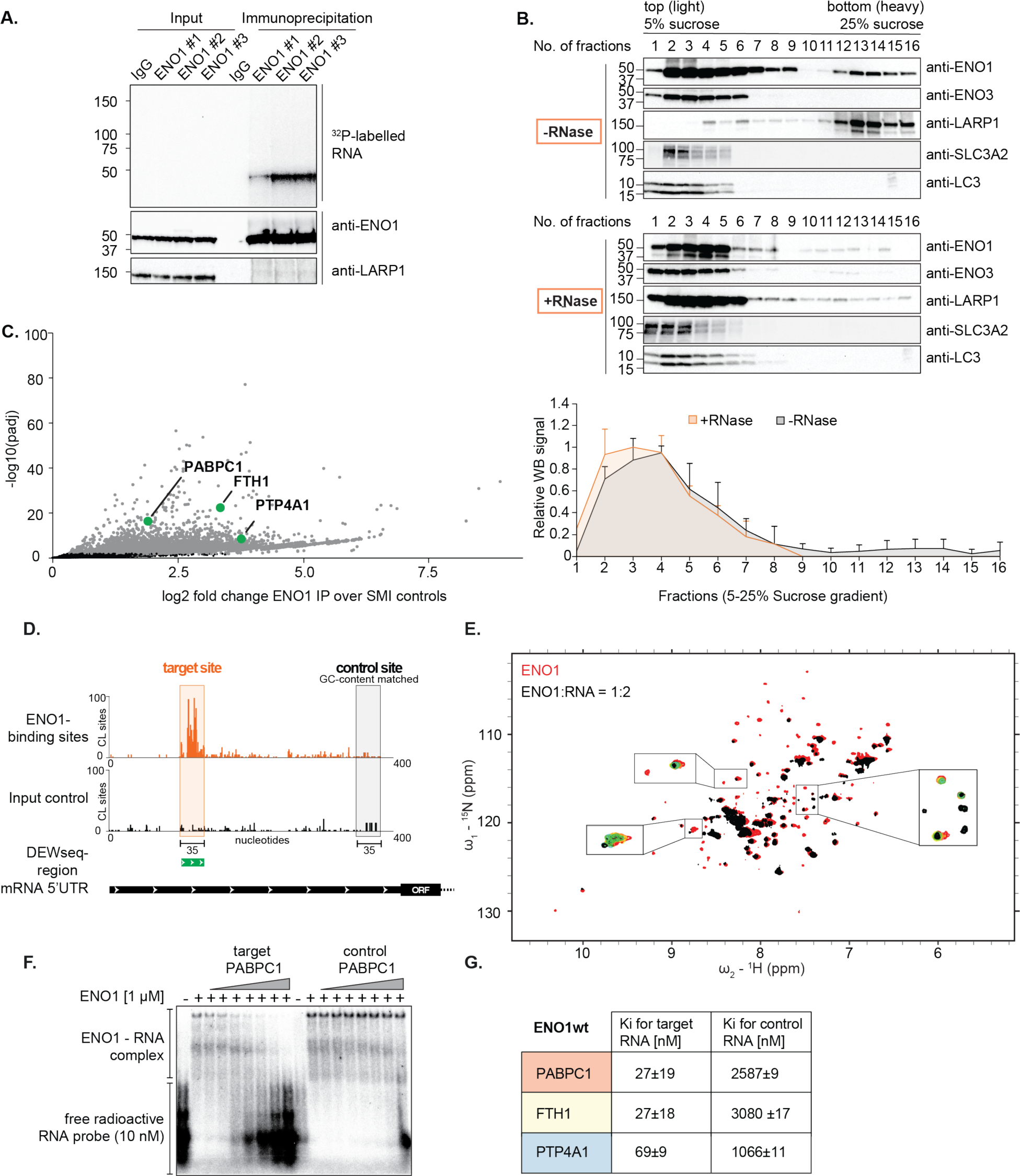
The glycolytic enzyme Enolase 1 binds specific RNAs in vivo and in vitro. A. ENO1 immunoprecipitation (IP) after crosslinking RNA-RBP complexes with UV light. Polynucleotide kinase labels RNA with radioactive ^32^P-yATP (PNK assay). Western blot staining for ENO1 and LARP1 of the same experiment. Three biological replicates are shown for ENO1. B. Top: HeLa cell lysates were treated with RNase I/A/T1 or left untreated, subjected to sucrose density gradient (5–25%), centrifugation and fractionation. A protein’s RNase-sensitive fraction moves from the higher fraction number (heavy) to the lower fraction number (light). Bottom: Quantification of biological replicates for the experimental setup on the top for ENO1 immunoblots (n=3, SD is indicated). C. Volcano plot of differential crosslinking site occurrences as determined by DEWseq; each dot corresponds to a window of the genomic region (50 nucleotides), grey coloring indicates significant enrichment in ENO1 IPs (IHW-adjusted p-value <0.1, log2 FC >0.5). The data are based on five biological experiments and were normalized to the background. Only the positive enrichment is displayed. Green circles indicate ENO1 target sites selected for further experiments. D. Schematic of ENO1’s binding profile as identified by eCLIP relative to the input control used for the identification of target and control sites. Every line represents an accumulation of crosslink sites at an individual nucleotide. E. Overlay of ^1^H, ^15^N-HSQC spectra of free ENO1 (red) and ENO1 incubated with twofold excess of RNA (FTH1 18-mer start, black; Fig. S2). Full titration points (in 1:0.1/0.2/0.4/0.8/1.2/2 ratios) of some residues are shown in insets. F. Comparative electromobility shift assay (EMSA) for target versus control RNA using radioactively labelled PABPC1 target 35-mer as a probe and unlabeled competitor RNA from either the target or control region as indicated in A. G. Inhibition constants (Ki) for target and control RNAs for the binding to ENO1wt (n=3). The Ki was calculated using a non-linear fitting with least squares regression.

We next determined the RNA-binding sites of ENO1 in the transcriptome by applying the enhanced crosslinking and immunoprecipitation (eCLIP) protocol (*13*)(Fig. S1A–C). We attained that ENO1 interacts with a wide range of RNAs in HeLa cells with a preference towards the 5’untranslated region (5’UTR) of mRNAs (Fig. S1D). Based on the exact crosslinking sites and our DEWseq analysis tool (*14*), we identified approximately two thousand direct ENO1-binding sites across the transcriptome (Fig. 1C) that do not share a striking linear sequence motif signature. The top scoring two sequence motifs (Fig. S1E) jointly account for only ~22% of all ENO1 binding sites that we identified.

For validation experiments, we synthesized RNAs of 35 nucleotides in length that either correspond to ENO1’s binding site or a GC content-matched control, derived from a region of the same mRNA downstream from its binding site (schematic in Fig. 1D). We tested an exemplary ligand and control RNA, derived from the PABPC1 5’UTR in a competition electromobility shift assay (EMSA) using recombinant human ENO1 (Fig. 1F and G; Ki_target_: 27±19 nM; Ki_control_: 2587±9 nM, Fig. S2B). Similar results were obtained for two additional ligand and control pairs derived from the PTP4A1 and FTH1 mRNAs, respectively (Fig. 1G, Fig. S2C and D). Using NMR, we observed RNA-induced chemical shift perturbations and line broadening of ENO1 resonances in ^1^H, ^15^N-HSQC spectra, confirming direct RNA binding in vitro (shortened FTH1 ligand RNA (18-mer), Fig. 1E, Fig. S2A). Taken together, ENO1 binds RNA at numerous transcriptomic sites in human cells with two orders of magnitude difference between specific and non-specific interactions.

To uncover the functional relevance of the interaction between ENO1 and RNA, we knocked down endogenous ENO1 and collected RNAseq data to identify the effect of ENO1 on its RNA targets. Despite an efficient knock-down of ENO1 (Fig. S2F), we could not obtain convincing evidence for a regulatory effect of ENO1 on the transcriptome in general or the ENO1 targets determined by eCLIP (Fig. S2E, G and H). Motivated by recent data on the regulation of p62 function in autophagy by RNA (*15*) and classic work on the control of E. coli RNA polymerase by 6S RNA (*16*), we tested whether conversely RNA binding might affect ENO1’s enzymatic activity. To this end, we purified recombinant human ENO1 and assayed its activity in the presence of RNA (Fig. 2A and B, Fig. S3A and B). While the three, independent ligand RNAs inhibit ENO1’s activity reproducibly and to a similar extent (Fig. 2A and B, Ki_PABPC1_: 99.1 nM, 95% CI: 49.42–223.2 nM; Fig. S3A, Ki_FTH1_: 150 nM, 95% CI: 86.4–310.9 nM; Fig. S3B, Ki_PTP4A1_: 103 nM, 95% CI: 56.0–190.3 nM), the corresponding control RNAs do not elicit significant changes in ENO1’s activity. Modelling of the data suggests a non-competitive inhibition (*17*) mode for the interaction between ENO1 and RNA.

**Fig. 2.**
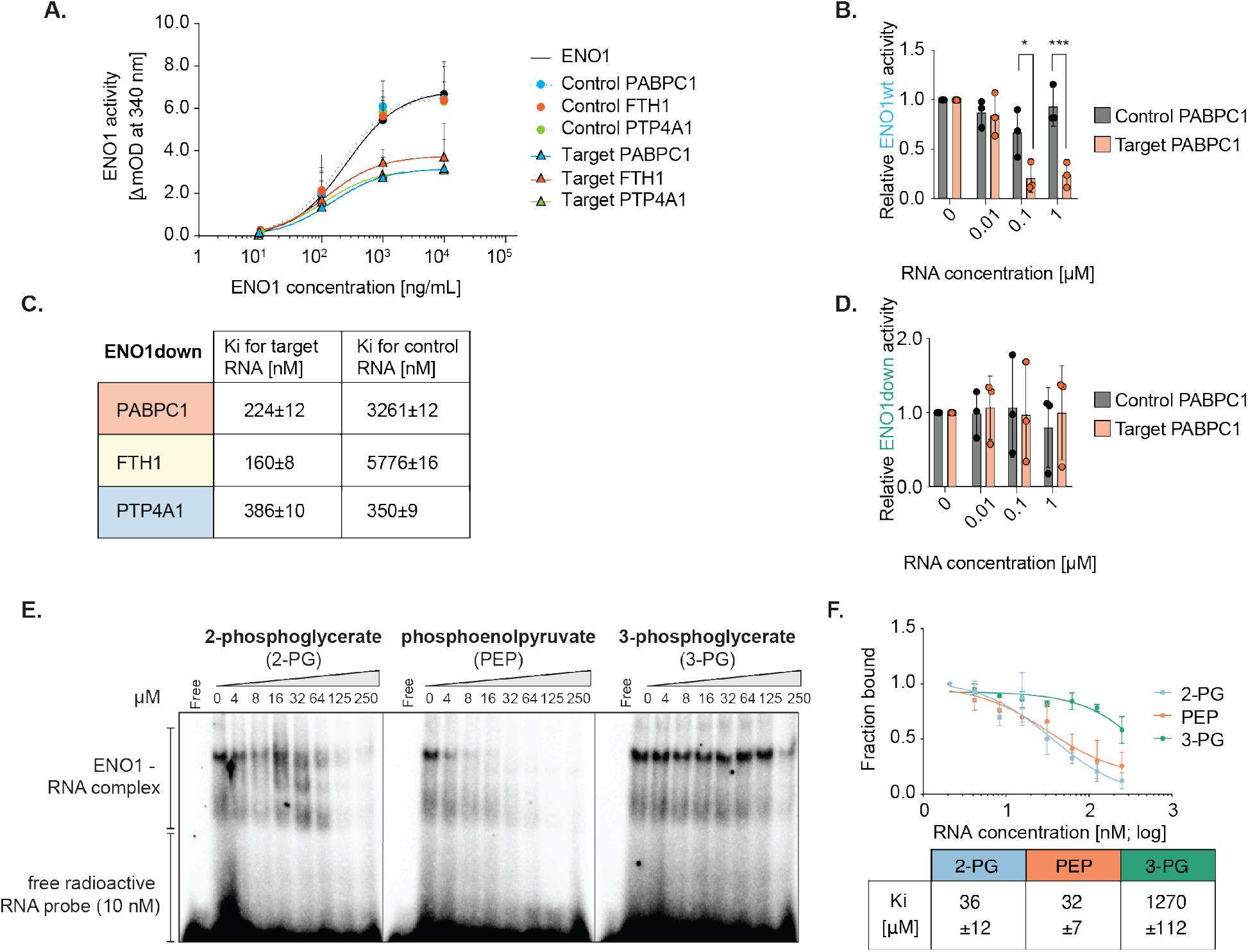
Riboregulation of ENO1’s enzymatic activity in vitro. A. Enzymatic activity assay of recombinant human ENO1 with increasing enzyme concentrations exposed to different control and target RNAs (constant conc. 100 nM). B. Measurement of recombinant ENO1wt activity with increasing concentrations of control and target PABPC1 RNA (n=3, SD is indicated). Statistically significant differences were detected using two-way ANOVA and Sidak-correction for multiple comparison testing. C. Inhibition constants (Ki) for target and control RNAs for the binding to ENO1down determined by competitive EMSA (n=3). The Ki was calculated using a non-linear fitting with least squares regression. D. Measurement of recombinant ENO1down activity with increasing concentrations of control and target PABPC1 RNA (n=3). E. EMSA with recombinant human ENO1 and labelled PABPC1 target RNA in competition with substrate of the forward (2-phosphoglycerate) and reverse reaction (phosphoenol pyruvate) or a control glycolytic metabolite (3-phosphoglycerate). F. Quantification of replicates for the experimental setup in E, including the Ki (n=3). The Ki was calculated using a non-linear fitting with least squares regression.

Instructed by RBDmap data (*15, 18*), we generated an ENO1 mutant (K343A; ENO1down) with ~5–10-fold decreased RNA binding (Fig. 2C and D; Ki_target_: 224±12 nM; Ki_control_: 3261±12 nM, Fig. S3C–E) compared to ENO1wt as measured by competitive EMSA (Fig. 1F and G, Fig. S2B–D). Importantly, ENO1down displays no discernible alteration in its enzymatic activity with any of the RNAs tested (Fig. 2D, Fig. S3F and G). Thus, both control RNAs and the ENO1down mutant corroborate that RNA specifically riboregulates ENO1’s activity in vitro.

Next, we tested whether ENO1’s enzymatic substrate binding and RNA binding are competitive. For this reason, we performed EMSA experiments utilizing 2-PG and PEP (*19*) as competitors. While both substrates compete with RNA for ENO1’s binding, the specificity control 3-phosphoglycerate (3-PG), the immediate precursor of 2-PG with an identical molecular mass fails to compete (Fig. 2E and F). Thus, we observe specific competition between substrates and RNA ligands for binding to ENO1.

In addition to ENO1down, we generated the ENO1up mutant. The design of ENO1up was also guided by RBDmap data (*18*) and entails the change of lysine residues (K89A/K92A/K105A) along the inferred interaction region of RNA with the enzyme. After knocking down endogenous HeLa cell ENO1 and rescue with the respective Flag-tagged ENO1 variant, ENO1up displays increased RNA binding compared to ENO1wt, as measured by PNK assays (Fig. 3A and B). In line with the in vitro data (Fig. 2C, Fig. S3C–E), ENO1down displays substantially decreased RNA binding in HeLa cells relative to ENO1wt (Fig. 3A).

**Fig. 3.**
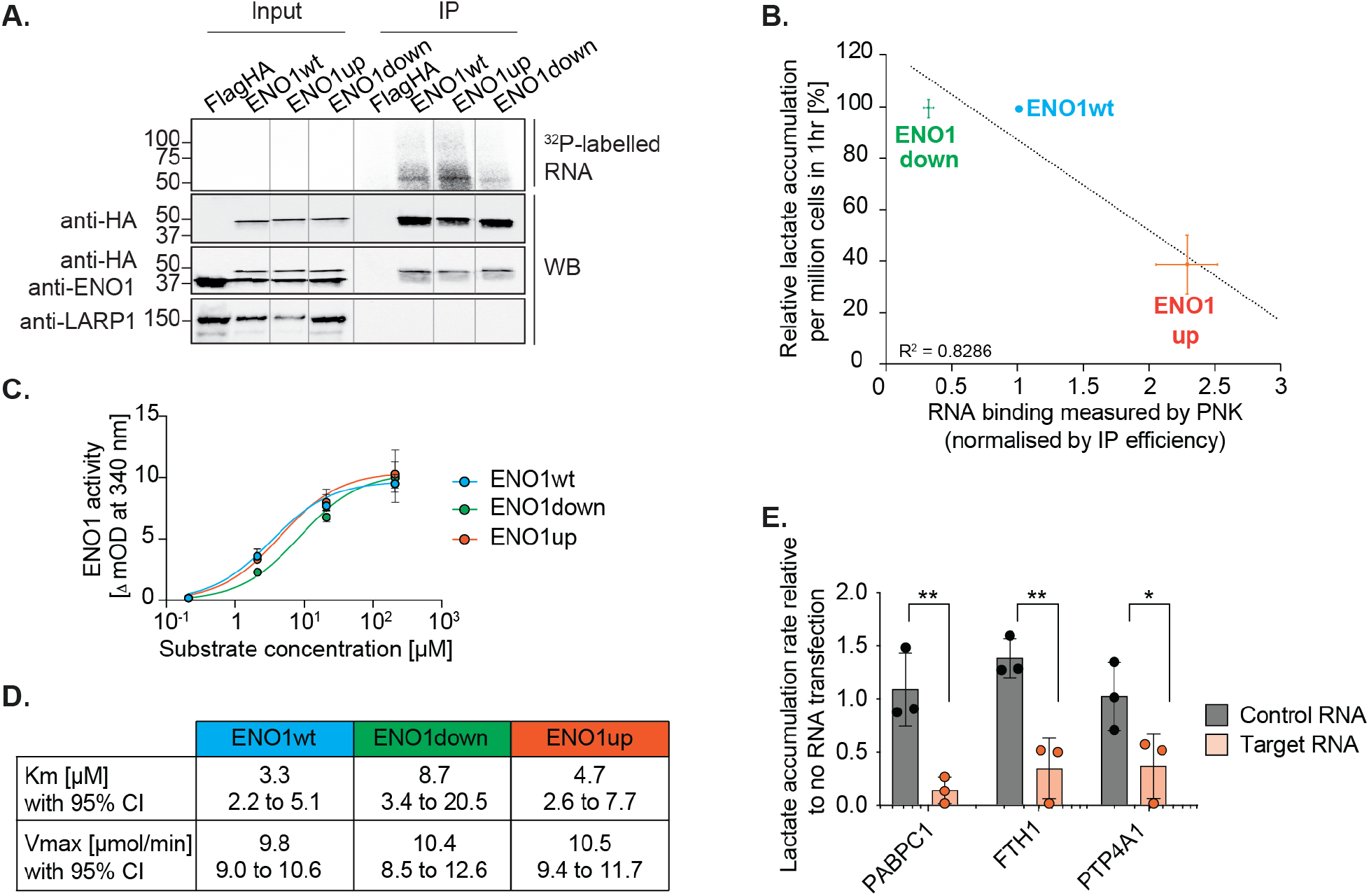
Riboregulation of ENO1 in HeLa cells and mESCs. A. Flag immunoprecipitation and PNK assay of transiently expressed ENO1-Flag-HA proteins after knocking down endogenous ENO1 with siRNAs. RNA binding of wild-type ENO1 (wt), a mutant with increased RNA binding (ENO1up) and reduced RNA binding (ENO1down) are being compared. B. Comparison of ENO1’s RNA-binding (PNK), and ENO1’s enzymatic activity in HeLa cells, indirectly measured by the accumulation of lactate in the medium (n=3). C. Michaelis-Menten saturation curve of the basal enzymatic activity of recombinant ENO1wt, ENO1down and ENO1up in vitro in the absence of RNA using a non-linear curve fitting with least squares regression (n=3). D. Vmax and Km measurements for ENO1wt, ENO1down and ENO1up as determined from the Michaelis-Menten saturation curve in c. The asymmetrical confidence interval (CI) is given (n=3). E. Three distinct sets of target and control RNAs (5 μM) were nucleofected into mESCs. Upon nucleofection of the control or target RNAs, the lactate accumulation in the medium was measured after 30, 60 and 90 minutes and used to estimate the accumulation rate by calculating the slope. The R^2^ value was used as a quality control. The two-tailed Student’s *t*-test is used to detect statistically significant differences (n=3, SD is indicated).

When we tested these mutants for their ability to rescue glycolysis (lactate accumulation in the medium) in HeLa cells after knock-down of the endogenous ENO1, ENO1wt expectedly rescued lactate production (Fig. 3B). Of note, the RNA binding-deficient mutant ENO1down is as active as the wild-type protein (Fig. 3B), showing that the K343A mutation does not incapacitate the enzyme. In contrast, ENO1up fails to rescue the knock-down-induced inhibition of lactate accumulation (Fig. 3B), although it is fully active when tested in the absence of RNA in vitro (Fig. 3C and D, Fig. S4A). The activity measurements were controlled with a mutant lacking enzymatic activity (ENO1as, E295A/D320A/K394A, Fig. S4A)(*20*). These results are consistent with the notion that EN01’s RNA binding interferes with its enzymatic activity in cells. We complemented these experiments by nucleofection of ligand and control synthetic 35-mer RNAs into HeLa cells. The results unambiguously confirm specific riboregulation of lactate production by the ENO1 ligand RNAs tested (Fig. S4B and C).

To explore physiological functions of the ENO1-RNA interaction, we chose mouse embryonic stem cells (mESCs). Like many cancer cells, mESCs utilizes glucose as a major energy source in the undifferentiated state (*1, 2*). Removal of the leukaemia inhibitory factor (LIF) from the culture medium induces differentiation, accompanied by a decrease in glycolysis and increased respiration (Fig. S5A and B). Interestingly, the decrease in glycolysis correlates with increased ENO1 RNA binding after LIF withdrawal (Fig. S5C).

To directly test the effect of RNA on ENO1 in mESCs, we nucleofected control or ligand RNAs and measured lactate accumulation in the medium. Confirming the results from HeLa cells (Fig. S4B and C), all three specific ENO1-binding RNAs significantly reduce lactate accumulation, in stark contrast to the control RNAs (Fig. 3E).

Unfortunately, the RNA nucleofection protocol is incompatible with meaningful mESC differentiation analyses. For this reason, we used unperturbed mESCs (*21*), withdrew LIF for a period of seven days, and sorted cells that were positive for the expression of Brachyury (BFP-positive), which is primarily found in cells differentiating towards the primitive streak (*22*), or Eomes (mCherry-positive), which is predominantly expressed in the definitive endoderm (*23*). We detected that lactate accumulation in the medium of Eomes+ cells significantly exceeds that of Brachyury+ cells (Fig. 4A), suggesting that the differentiation to the definitive endoderm may require sustained glycolysis in comparison to the primitive streak. Of note, ENO1 binding to RNA correlates inversely (compare Fig. 4A and B).

**Fig. 4.**
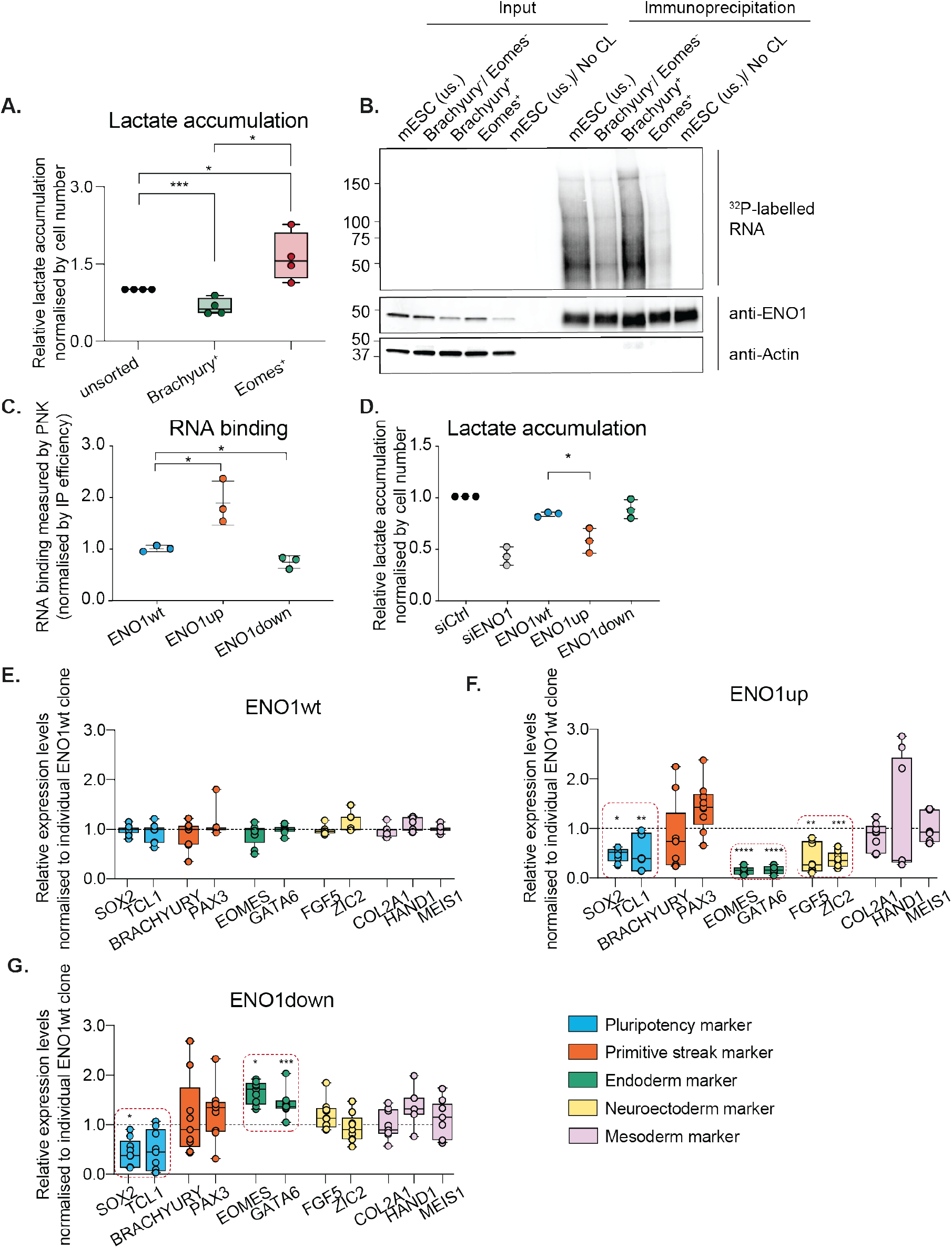
Riboregulation of ENO1 affects mESC differentiation. A. Cells expressing Eomes-mCherry or Brachyury-BFP after seven days of LIF withdrawal were sorted for the respective fluorescent marker proteins, cultured for an additional five days and lactate accumulation in the culture medium was quantified (n=3, SD is indicated). The statistically significant differences were detected using the two-tailed Student’s *t*-test. B. Cellular RNA-protein interactions were cross-linked with UV and a PNK assay was performed for cell populations described in A. Non-crosslinked (No CL), unsorted (us.) mESC were used as specificity controls. Immunoblots of ENO1 and actin are shown as controls. C. Cell lines were established for ENO1 knockout and constitutive transgenic expression of Flag-HA-tagged ENO1wt, ENO1up and ENO1down from the Rosa26 locus. RNA binding was measured for the three ENO1 variants by Flag immunoprecipitation and PNK assay after UV cross-linking in pluripotent mESCs. Three independent clones were used. The standard deviation is given and the statistically significant differences were detected using the two-tailed Student’s *t*-test when comparing to ENO1wt. D. Measurement of the lactate accumulation in the culture medium for three independent clones of ENO1wt, ENO1up and ENO1down in pluripotent mESCs. The standard deviation is given and the statistically significant differences were detected using the two-tailed Student’s *t*-test when comparing to ENO1wt. E.–G. Differentiation of mESCs and qPCRs of lineage marker for ENO1wt (E.), ENO1up (F.) and ENO1down (G.) after seven days of LIF withdrawal for three independent clones (from C.) in three experiments. The standard deviation is given and the statistically significant differences were detected using two-way ANOVA with Tukey-corrected multiple comparison testing when comparing to ENO1wt.

To examine whether this correlation reflects a causal requirement for riboregulation of ENO1 during ESC differentiation, we knocked out endogenous ENO1 and introduced murine versions of the ENO1 variants, characterized above, into the Rosa26 locus (Fig. 3A) by CRISPR/Cas9 genome editing. The different heterologous forms of ENO1 are expressed at similar levels and below the expression level of endogenous ENO1 (Fig. S5D). As one would expect, lactate accumulation in the medium of ENO1wt cells is somewhat less than seen in control cells (Fig. 4D). As previously observed in HeLa cells, ENO1up displays increased RNA binding and mediates decreased lactate production; likewise, ENO1down shows a decrease in RNA binding compared to ENO1wt (Fig. 4C and D).

Independent clones of these cell lines were subjected to LIF withdrawal and analyzed for differentiation to the different germ layers. Engineered mESCs expressing ENO1wt differentiated normally into the distinct germ layers, as assessed by qPCR analysis of the respective expression of marker genes (Fig. 4E). By contrast, ENO1up-expressing cells fail profoundly in their differentiation to definitive endoderm and neuroectoderm (Fig. 4F), while the expression of primitive streak and mesodermal markers was quite variable and statistically not significantly affected. We also noticed that ENO1down cells, where ENO1’s activity escapes riboregulation, show increased differentiation towards the definitive endoderm (Fig. 4G).

Taken together, these results demonstrate physiological ENO1 riboregulation in the course of mESC differentiation and its requirement especially for the formation of the endodermal germ layer. As such, these results represent a novel form of regulated cell differentiation.

Our biochemical studies and the data using the ENO1 mutants show that RNA directly interacts with the enzyme in a way that is mutually exclusive with substrate binding. Based on RBDmap, we surmise that ENO1down is RNA-binding deficient because K343 directly interacts with RNA; this hypothesis awaits structural elucidation. The finding that many, possibly thousands of sites within mammalian cell transcriptomes display specific ENO1 binding, as exemplified by the PABPC1, PTP4A1 and FTH1 target sites, raises the question of how regulation is achieved, e.g. during mESC differentiation. Although this question remains to be answered experimentally, a posttranslational modification that controls the overall ability of ENO1 to bind RNA represents an attractive possibility. The ENO1up mutant, as identified in RBDmap and RBS-ID data (*24*), suggests candidate sites for such a modification. According to this scenario, the transcriptome as a whole bearing thousands of relevant ENO1-binding regions would serve a specific regulatory function, a concept that not many will consider obvious. Our findings could pave the way towards a novel class of compounds that hijack the cells’ endogenous riboregulatory mechanisms for therapeutic intervention (*25*).

## Acknowledgements

We thank S. Sahadevan for his expert support with the data analysis and M. Büscher for advice on the knock-out and knock-in experiments. We also thank current and former members of the Hentze laboratory for their feedback, K-M. Noh (EMBL) for sharing the pSpCas9(BB)-2A-RFP plasmid and D. Kuster for technical help. We acknowledge EMBL’s core facilities, specifically the Genomics, Flow Cytometry and Metabolomics Core Facilities for their expert services.

## Author Contributions

I.H. contributed to conceptualization, formal analysis, investigation, methodology, project administration, visualization and writing (original draft and editing); J. I. P.-P. contributed to investigation and methodology (mESCs); P. M., T. S. and T. S. conducted the bioinformatical analyses (eCLIP, RNAseq); L. D.-P. and J. H. contributed to the acquisition, analysis and the interpretation of data (NMR); P. N. contributed to the design and interpretation of the work, M. W. H. contributed to conceptualization, data analysis and interpretation, funding acquisition, project administration, mentoring and writing (original draft and editing).

## Data availability

The principal data supporting the findings of this Article are available within the figures and the Supplementary Information; eCLIP (E-MTAB-9031) and RNAseq data (E-MTAB-9032) generated for this study were deposited in Array Express.

## Supplementary Materials

### Materials and Methods

#### Cell culture and transfection

HeLa cells (human female origin) were cultured in high-glucose (10 mM) Dulbecco’s Modified Eagle Medium (DMEM), supplemented with 10% heat-inactivated fetal bovine serum (FBS, Merck), 2 mM L-glutamine (25030081, Thermo Fisher) and 100 U/ml Pen/Strep (15140122, Thermo Fisher). For the transfection of HeLa cells with siRNAs, cells were grown until 70%–80% confluent, trypsinized, counted (TC20, Bio-Rad) and 200,000 cells were seeded per well in a 6-well dish for reverse transfection of siRNAs following the manufacturer’s instructions (137780755, Lipofectamine RNAiMax, Invitrogen). For the transfection of plasmids, 5 μL of Lipofectamine 2000 reagent were used combined with 1 μg of plasmid DNA. In case of combined knock-down and rescue experiments, the cells were seeded and siRNA transfection was performed (ENO1 pool: SMARTpool ON-TARGETplus L-004034-00-0005 and control pool: ON-TARGETplus non-targeting siRNAs 1–4). After 24 hours, the plasmid was transfected followed by an additional 24-hour incubation. The mouse embryonic stem cells (R1) were a donation from the group of Alexander Aulehla. Cell culture dishes were coated with 0.1% gelatine (G1890, Sigma) in PBS for 15 minutes prior to the addition of mESCs. The mESCs were cultured in high-glucose (10 mM) DMEM (D5796, Merck), supplemented with 15% FBS (EmbryoMax, ES-009-B, Merck Millipore), 2 mM L-glutamine (25030081, Thermo Fisher), 100 U/ml Pen/Strep (15140122, Thermo Fisher), 100 μM MEM non-essential amino acid solution (11140035, Merck), 1 mM sodium pyruvate (11360088, Merck) and 0.1 mM β-mercaptoethanol. If the mESCs were maintained to remain pluripotent, the leukaemia inhibitory factor (10^3^ units/ml) was added to the medium. The triple knock-in cell line was a donation from the group of Pierre Neveu (*21*) and was cultured under the same conditions. The cell lines were not authenticated.

#### CRISPR/Cas9 gene insertion of ENO1 mutants

Single guide RNAs targeting the ROSA locus were predicted using the CRISPR online tool (*26, 27*), ordered from Sigma-Aldrich, annealed and cloned into the pSpCas9(BB)-2A-RFP (kindly provided by Kyung-Min Noh, EMBL) using the BbsI restriction sites (kindly cloned by David Kuster). The template plasmids for the knock-in of ENO1 mutants were prepared by introducing homology arms complementary to the ROSA locus upstream and downstream of the ENO1 wildtype or mutant DNA. While ENO1 is expressed under an EF1alpha promoter, the fluorescent protein cerulean is expressed under an additional, separate IRES promoter to facilitate an efficient cell sort. The generated plasmids were nucleofected into mESCs using the Nucleofector 4D system according to the manufacturer’s guidelines (Lonza, Cell Line Nucleofector Kit P3, program CG-104, one million cells, 16 μg template vector and 4 μg Cas9 vector per nucleofection). Single cell sorting of double positive cells (RFP and cerulean) was performed 48 hours after nucleofection. Upon clonal expansion, successful insertion was tested by PCR to screen for homozygous insertions of the ENO1 wildtype or mutant DNA sequence at the ROSA locus (Phire Tissue Direct PCR Master Mix, F170L, Takara). The mutants of ENO1 were based on the findings of RBDmap data (*15, 18*).

#### CRISPR/Cas9 gene deletion of ENO1

Using the ENO1 wildtype or mutant-expressing cell lines, we knocked out ENO1 in a subsequent step. Guide RNAs targeting the ENO1 locus were predicted using the CRISPR online tool (*26, 27*), ordered from Sigma-Aldrich, annealed and cloned into the pSpCas9(BB)-2A-GFP (PX458) (Addgene plasmid no. 48138, kindly provided by Fang Zhang), or pSpCas9(BB)-2A-RFP (kindly provided by Kyung-Min Noh, EMBL) using the BbsI restriction sites. The generated plasmids were nucleofected into mESCs using the Nucleofector 4D system according to the manufacturer’s guidelines (Lonza, Cell Line Nucleofector Kit P3, program CG-104, 1 million cells and 5 μg of each of the guide RNA-containing plasmids). Single cell sorting of double positive cells (RFP and GFP) was performed 48 hours after nucleofection. Upon clonal expansion, successful insertion was tested by PCR to screen for homozygous deletion of ENO1 (Phire Tissue Direct PCR Master Mix, F170L, Takara). Furthermore, we tested the expression of ENO1 on a Western blot.

#### Differentiation and Fluorescence-activated cell sorting

To induce differentiation, LIF was removed from the medium for a period of seven days before harvesting (15 cm dishes). The cells were then trypsinized using 5 ml TrypLE Express at 37°C for five minutes and sorted using a BD Aria Fusion Sorter. Cell populations of BFP-positive, GFP-positive and mCherry-positive cells were respectively plated on gelatine-coated 10 cm dishes using LIF-free medium and cultured for five additional days before performing further experiments.

#### Protein extracts, SDS-PAGE and Western blotting

For Western blotting, cells were washed twice with ice-cold PBS and lysed on the plate using RIPA lysis buffer, supplemented with protease inhibitor (11873580001, Roche). Lysates were treated with benzonase (100 U/ml, 71206, Merck Millipore) for 15 minutes on ice and the protein concentrations were measured using the Bio-Rad Protein Assay. 4× loading buffer was added to the lysates, boiled for five minutes and typically 15 μg of lysate was used for SDS-PAGE. Proteins were transferred to nitrocellulose membranes using the Trans-Blot Turbo Transfer System (Bio-Rad) and blocked for 1 hour at room temperature with 5% milk in TBST. Primary antibodies were incubated in 5% milk TBST either overnight at 4°C or one hour at room temperature (RT), followed by 3× TBST washes, secondary antibody incubation in 5% milk in TBST for one hour at RT, 3× TBST washes and developed using ECL (WBKLS0500, Millipore). The antibodies used for Western blotting are listed here: anti-ENO1 (1:10,000; 11204-1-AP, Proteintech; 1:5,000; ab112994, Abcam), anti-ENO3(1:2,000, H00002027-M01, Novus), anti-LARP1 (1:2,000; A302-088A, Biomol), anti-SLC3A2 (sc-9160, Santa Cruz), anti-LC3B 1:300 (CTB-LC3-2-IC, Cosmo Bio), β-actin (1:5,000; A5441, Sigma-Aldrich) and HA (1:3,000; 901502, BioLegend). Secondary antibodies (goat anti-mouse IgG-HRP, ab97023; goat anti-mouse IgG-HRP, ab6721; rabbit antigoat IgG-HRP; ab6741) were used at a 1:5,000 dilution. In the case of immunoprecipitations, the following secondary antibodies were used instead due to the overlap in size of ENO1 and the IgG heavy chain (anti-rabbit light chain HRP antibody, MAB201P, Millipore; anti-mouse light chain, HRP antibody, AP200P, Millipore).

#### Polynucleotide kinase assay

Prior to harvesting cells, ENO1 antibody (ab112994 for HeLa cells or ab15595 mouse embryonic stem cells, Abcam) was coupled with Protein G magnetic beads (SureBeads, 161-4023, Bio-Rad) for one hour at room temperature. Typically, 1 μg was coupled with 30 μl of bead slurry. After coupling, the magnetic beads were washed twice with lysis buffer (100 mM NaCl; 50 mM Tris-HCl pH 7.5; 0.1% SDS; 1 mM MgCl_2_; 0.1 mM CaCl_2_; 1% NP-40; 0.5% sodium deoxycholate; protease inhibitors (11873580001, Roche)). Alternatively, pre-coupled Flag M2 magnetic beads (M8823, Sigma Aldrich) were washed twice with lysis buffer (25 μL per immunoprecipitation). HeLa cells or mESCs were washed twice with ice-cold PBS. Subsequently, the cells were UV-crosslinked at 150 mJ/cm^2^ on ice and lysed in lysis buffer. The lysates were sonicated (BioruptorPlus) and treated with 10 U/ml RNase I (AM2295, Thermo Fisher) and 2 U/ml Turbo DNase (AM2238, Thermo Fisher) for five minutes at 37°C. The lysates were centrifuged full speed at 4°C for 15 minutes and 15 μl were saved as inputs for a Western blot. The remainder of the lysates was then utilized for immunoprecipitations (IPs) at 4°C for two hours. After the IP and three times three-minute washes with lysis buffer at room temperature, beads were washed additionally twice with PNK buffer (50 mM NaCl; 50 mM Tris-HCl, pH 7.5; 10 mM MgCl_2_; 0.5% NP-40; protease inhibitors (11873580001, Roche)). The magnetic beads were resuspended in PNK buffer containing 0.1 mCi/ml [γ-^32^P] ATP (Hartmann), 1 U/ml T4 PNK (NEB), 1mM DTT and labelled for 15 minutes at 37°C. After four washes with PNK buffer, proteins were eluted at a low pH (0.1 M glycine, pH 2.0), neutralized with 1.5 M Tris-HCl, pH 8.5, and mixed with 4× sample loading buffer. The samples were heated to 95°C for five minutes and quickly spun down. The samples were resolved by SDS-PAGE and blotted on a nitrocellulose membrane using the Trans-Blot Turbo Transfer System. The membrane was exposed overnight to a phosphorimaging screen, followed by immunoblotting.

#### eCLIP

eCLIP was performed as previously published (*13*), with minor adaptations. 1-3 μg of ENO1 antibody (ab112994, Abcam) or appropriate control IgG (rabbit) was coupled for 1 hour at RT to 30 μl of Protein G coupled magnetic beads. The cell lysates were treated with 0.1 U/ml RNase I (AM2295, Thermo Fisher) for five minutes. One ml of lysate was used for each immunoprecipitation (concentration of 2 mg/ml) for two hours at 4°C. Complexes were eluted at low pH (0.1 M glycine, pH 2.0) and neutralized with 1.5 M Tris-HCl pH 8.5 before loading them on Criterion XT Bis-Tris Protein Gels (3450124, Bio-Rad). The transfer onto the nitrocellulose membrane (1704271, Bio-Rad) was performed using the Trans-Blot Turbo Transfer System. cDNA libraries composed of the IP (16 cycles) and the size-matched input controls (9 cycles) were multiplexed and sequenced using paired-end sequencing (PE125) on the Illumina HiSeq2000 platform.

#### Data processing for eCLIP and statistical analysis

The quality check of the eCLIP data was performed using fastqc (*28*). The Unique Molecular Identifier (UMI) barcodes that are attached during library preparation were extracted and appended to the read name using umi-tools extract (*29*). The ligated adapters were trimmed and the reads shorter than 18 nucleotides were discarded using the cutadapt tool (*30*). The adapter-trimmed reads were aligned to the human genome (GRCh38.v23 from GENCODE) with STAR (*31*), a splice-aware aligner, with the end-to-end alignment mode. The PCR duplicated reads were filtered from the mapped reads, based on the UMI barcodes appended to the read name, using umi-tools dedup with default parameters.

The GENCODE annotation of GRCh38.v23 genome was extended by including coordinates for tRNAs provided by tRNAscan (*32*), and the resulting annotation was pre-processed with the htseq-clip suite (available at https://htseq-clip.readthedocs.io/en/latest/overview.html; accessed 21 May 2020) into overlapping windows of 50 nucleotides in size with a step size of 20 nucleotides. The truncation site (position-1 relative to the start site of a read, also called crosslink site) was extracted and quantified using htseq-clip. We used the R/Bioconductor DEWSeq package (*14*) to detect significantly enriched windows in immunoprecipitation samples over the corresponding size-matched input control samples (log2 fold change >0.5, p-adjusted <0.1). IHW (*33*) was used for multiple hypothesis correction. Overlapping significant windows were merged to binding regions.

#### ENO1 Motif analysis

The previously obtained significant binding regions of ENO1 were filtered to retain regions with a minimum average of 30 normalized cross-link counts in the immunoprecipitation samples of protein-coding genes. We retained 472 protein coding exon regions, which were used for motif analysis. DREME(v4.11.3) (*34*) from the MEME suite package (*35*) was used to identify the de novo motifs with parameters maxk set to 20 and norc.

#### RT-PCR and qPCR

1.0 μg total RNA was used to synthesize cDNA using the Maxima First Strand cDNA Synthesis Kit (K1672, Thermo Fisher) for quantitative reverse transcription polymerase chain reaction (RT-qPCR). RT-qPCR was performed using the SYBR-Green qPCR Master mix (4309155, Applied Biosystems) on a StepOne Real-Time PCR System (Applied Biosystems, 384-well plate). Gene expression values were normalized using ACTB and are shown as a relative fold change to the value of control samples. All experiments were performed in biological triplicates and error bars indicate ± standard deviation as assayed by the ΔΔCt method. Statistically significant differences were detected using two-way ANOVA with multiple comparisons correction (Tukey). All RT-qPCR primers are listed in Supplementary Table 1.

#### RNAseq library preparation

RNA was isolated using the Direct-zol RNA Miniprep kit (R2051, Zymogen), as recommended by the manufacturer. 1.0 μg total RNA was used as input for the library preparation. The TruSeq Stranded mRNA kit was used for the library preparation. The samples were multiplexed and sequenced on a Hiseq2000.

#### RNAseq analysis and statistical analysis

Reads were trimmed using Cutadapt (*30*) (v1.16), mapped to the human genome (GRCh38.p3) with STAR (*31*) (v2.5.0.a) and summarized with featureCounts (*36*) (v1.6.4). Resulting counts were prefiltered to retain genes with more than 4 counts in at least 3 samples. DESeq2 (*37*) (v1.26) using local dispersion fit and likelihood ratio test with IHW (*33*) for multiple hypothesis correction was used to determine significantly enriched RNAs in ENO1 knock-down versus control samples (adjusted p-value <0.1; log2 fold change >0.5).

#### Sucrose density centrifugation and fractionation

For the preparation of lysates for the testing of RNA dependency through ultracentrifugation, previously published protocols were used as a basis (*38, 39*). HeLa cells were cultured on a 15 cm dish and lysed in 300 μl lysis buffer (25 mM Tris-HCl, pH 7.4, 150 mM KCl, 0.5% NP-40, 2 mM EDTA, 1 mM NaF, 0.5 mM DTT, protease inhibitor). A pre-clearing step was performed by centrifugation at 10,000g for 10 minutes at 4°C. The lysates were then treated with a combination of RNase I (100 U, AM2295, Thermo Fisher), RNase T1/A (100U, EN0551, Thermo Fisher) or no RNase was added at 37°C for 15 minutes. For the fractionation, gradients from 5% (w/v) to 25% (w/v) sucrose in 150 mM KCl, 25 mM Tris-HCl (pH 7.4) and 2 mM EDTA were prepared using a gradient maker (settings: 1:52, angle 81.5, speed: 15). Lysates were separated by centrifugation at 30,000 rpm and 4°C in an SW40 rotor for 18 hours. The lysate fractions were collected by hand through careful pipetting from the top (16 fractions were collected of approximately 600 μl). For the protein precipitation, 150 μl of 100% Trichloroacetic acid (TCA) was added and left on ice for 30 minutes. The individual fractions were centrifuged at full speed and 4°C for 20 minutes. The TCA supernatant was carefully removed and the pellets were washed once with 1 ml cold acetone (stored at −20°C). The samples were vortexed and an additional centrifugation step was performed at full speed and 4°C for 30 minutes. The supernatant was again carefully removed and the pellet was air-dried. Finally, the pellets were taken up in 1× loading buffer containing benzonase and used for SDS-PAGE and immunoblotting.

#### Protein Purification

The ENO1 wildtype and mutant DNA sequence was cloned into the pet22M plasmid (generously provided by the Protein Expression and Purification core facility at EMBL) using the NcoI and XhoI site. Through the expression of the ENO1 variants, the proteins are tagged with the protein Thioredoxin to add to the solubility of the protein, which can be cleaved off using the HRV 3C Protease and removed using a reverse Ni-NTA column.

For the protein purification, the ENO1 variants were expressed by transforming the plasmid into BL21 (DE3) competent cells and plated on Kanamycin-containing LB plates. One colony was picked and used for protein expression in a 400 ml flask. Once the OD600 of was reached, the culture was cooled down to 18°C and IPTG (I5502, Sigma Aldrich) was added to reach a final concentration of 1 mM. The bacteria were pelleted and lysed in lysis buffer (20 mM Tris-HCl pH 7.5, 500 mM NaCl, 5 mM b-2ME, 5% glycerol, 40 mM imidazole, 0.01% NP40), supplemented with RNase A, DNase and Protease Inhibitor to the lysis buffer. For efficient lysis, the lysates were processed with a microfluidizer. The His-tagged thioredoxin was used for the purification of ENO1 using a HisTrap HP column (17-5247-01, Amersham Biosciences) on an Äkta go protein purification system. The protein was eluted with increasing concentrations of imidazole and the protein-containing fractions were verified on a Coomassie gel. The solubility tag was cleaved by HRV 3C protease during overnight dialysis using the PUR tubing spectrapor 2. The following day, a reverse nickel column was performed using again a HisTrap HP column on an Äkta go protein purification system. The protein was eluted with the HRV 3C protease and Thioredoxin being retained due to their remaining His tag.

#### Electromobility shift assay (EMSA)

The synthesized RNA molecules (Sigma Aldrich) used for testing ENO1’s binding were radioactively end-labelled using the T4 PNK enzyme and [γ-^32^P] ATP (Hartmann). For all EMSA experiments, a 20 cm-long 5% acrylamide native gel was poured for the separation of the free radioactive probe and the formed complex of ENO1 and RNA. The reaction was performed in a 20 μl reaction mix, containing 1 μM ENO1 protein and 10 nM RNA in reaction buffer (1 mg/ml of BSA, 10 mg/ml RNAsin, 5 mM DTT, 0.5 mM PMSF, 2.5 mM MgCl_2_, 100 mM KCl; 20 mM HEPES, pH 7.9; 0.2 mM EDTA and 20% glycerol). Reactions were incubated at room temperature for 10 minutes. After the reaction samples were loaded on the native gel, it was run overnight at 55V. The next day, the gel was dried for 1 hour at 80°C and exposed to a phosphorimaging screen for four hours or overnight. The images were quantified using the ImageLab software and the modelling of binding curves of target and control RNAs was done using GraphPad Prism 8.

#### ENO1 activity assay

For measuring the activity of ENO1, the ENO1 activity assay kit (ab117994, Abcam) has been utilized according to the manufacturer’s instructions. For measuring the impact of RNA on the activity of ENO1. Synthesized RNAs (concentrations indicated in the respective figures) were added to the coupling step of the recombinant ENO1 to the ELISA plate. The plate was washed several times according to the manual and the reaction mix added. The non-competitive inhibition model was generated using GraphPad prism 8.

#### ENO1 Isotope labelling and protein purification

For the production of ^15^N-labelled, deuterated ENO1, BL21(DE3) cells expressing ecTRX-1-His6-tagged hsENO1 were pre-cultured in M9 minimal media supplemented with ^15^NH_4_Cl (Cambridge Isotope Laboratories) overnight at 37°C. The pre-culture was diluted in pre-warmed D_2_O (Cambridge Isotope Laboratories) M9 media to a final OD600 of 0.1. This second pre-culture was grown at 37°C to an OD600 of 0.8 and added to the final D_2_O M9 minimal media volume in a 1/10 dilution. The final culture was then grown to an OD600 of 0.9 at 37°C and hsENO1 expression was induced by adding 0.2 mM IPTG and grown overnight at 18°C.

The cells were harvested and resuspended in lysis buffer (20 mM Tris-HCl, pH 7.5, 500 mM NaCl, 40 mM imidazole, 5% glycerol, 0.01% NP-40, 5 mM β-mercaptoethanol, protease inhibitor cocktail (Roche)), followed by lysis with a microfluidizer (Microfluidics). The cleared lysate was applied to a 5 ml His trap FF column (GE Healthcare), washed and eluted with an imidazole buffer gradient (20 mM Tris-HCl, pH 7.5, 500 mM NaCl, 600 mM imidazole, 5% glycerol). The TRX-His6 tag was removed by addition of 3C PreScission protease. After dilution of the protein to lower the imidazole concentration (20 mM Tris-HCl, pH 7.5, 500 mM NaCl, 40 mM imidazole) the sample was applied to a second His trap column in order to remove the cleaved tag and the protease. The flow-through fraction was collected, concentrated and loaded onto a Superdex 200 16/600 size exclusion column (GE healtcare) equilibrated with NMR buffer (20 mM NaPi, pH 6, 150 mM NaCl, 20 mM MgCl_2_, 0.5 mM TCEP).

#### NMR data acquisition and titration

^1^H, ^15^N-HSQC spectra of deuterated ENO1 were recorded using an Avance III Bruker NMR spectrometer, which operates at a field strength, corresponding to a proton Larmor frequency of 800 MHz, equipped with a cryogenic triple resonance gradient probe head. Measurements were performed at 298 K, using 0.2 mM ENO1 in NMR buffer, supplemented with 5% D_2_O for the deuterium lock. This sample was titrated with increasing concentrations of 18-mer synthetic RNA (IBA) (0.02, 0.04, 0.05, 0.16, 0.24 and 0.4 mM) and at each titration step another ^1^H, ^15^N-HSQC was recorded. Data were processed and analysed using NMRPipe and Sparky.

#### Nucleofection of target and control RNAs

The synthesized specific and unspecific RNAs were nucleofected into mESCs using the Nucleofector 4D system according to the manufacturer’s guidelines (Lonza, Cell Line Nucleofector Kit P3, program CG-104, 1 million cells). The cells were then cultured for an additional 90 minutes in DMEM free of phenol red and FBS (D1145, Merck) before RNA extraction with the Direct-zol RNA Miniprep kit (R2051, Zymogen).

#### Lactate accumulation measurement

For the determination of lactate levels in the supernatant of HeLa and mESCs, the supernatant was collected 30, 60 and 90 minutes after the replacement with DMEM free of phenol red and FBS (D1145, Merck). The colorimetric assay kit for the determination of lactate levels was utilized according to the manufacturer’s instructions (K607, Biovision).

#### Lactate accumulation and oxygen consumption of pluripotent and differentiated cells

mESC were differentiated in 15 cm dishes for seven days in the absence of LIF or maintained for two days in the pluripotent state. Seeding of cells was done asynchronously so that pluripotent and differentiated cells could be processed in parallel on the same day. For determining oxygen consumption, cells were washed with PBS, trypsinized, pelleted by centrifugation (5 minutes at 230g) and resuspended in mESC medium (+ or - LIF as appropriate). Live cell number was counted in a TC20 automated cell counter (Bio-Rad) using trypan blue staining. Immediately afterwards, the oxygen consumption rate of 300 μl of cell suspension containing ~one million living cells was determined in an Oxytherm System (Hansatech Instruments) at 37°C using the Oxygraph Plus data acquisition software (Hansatech Instruments). For determining lactate secretion, the medium of the 15 cm dishes with pluripotent/differentiated cells was exchanged with 11 ml of mESC medium without serum and without phenol red. After 30 min of incubation, an aliquot of the conditioned medium was taken and stored at −80 °C. Cells were lysed in RIPA buffer and total protein quantified using the DC Protein Assay (Bio-Rad). Lactate concentration was measured in 3 μl of conditioned medium with the Lactate Assay kit (Bio-vision, catalogue K607) following manufacturer’s instructions. Lactate measurements were normalized by the protein amount on the dish.

#### Statistical analysis

Statistical analysis of results was performed using two-tailed Student’s t-test (parametric), or analysis of variance (ANOVA), followed by Sidak’s or Tukey’s multiple comparison tests, as stated in the figure legends. All the analyses were done using GraphPad Prism, version 8.4.2. Statistical significance is represented in all figures, as follows: p-value of <0.0001: ****, p-value of 0.0001 to 0.001: *** p-value of 0.001 to 0.01: **, p-value of 0.01 to 0.05: * and p-value of ≥0.05: not significant.

**Fig. S1.**
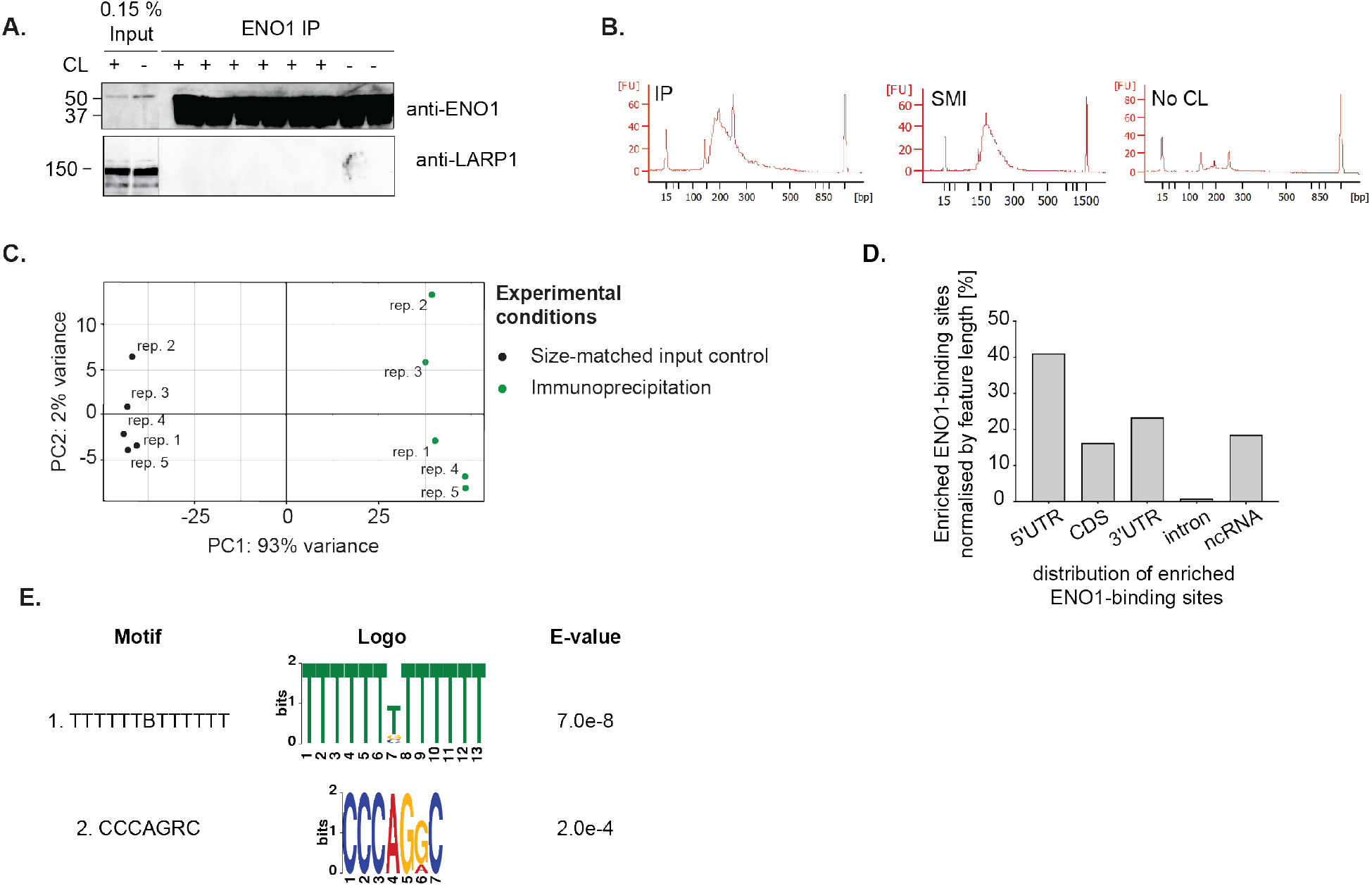
eCLIP analysis of ENO1 in HeLa cells. A. ENO1 immunoprecipitation (IP) efficiency after crosslinking RNA-RBP complexes with UV light. Six replicates of ENO1 eCLIP libraries were prepared. anti-LARP1 was stained as a control. B. Bioanalyzer traces of cDNA libraries for the ENO1 IP samples, the size-matched input (SMI) and the no crosslinking controls (No CL) C. Principal component (PC) analysis plot of ENO1’s eCLIP result comparing IPs and SMIs. One replicate (not shown) had to be excluded due to lower quality. D. Enriched ENO1-binding sites normalized by the length of the feature. E. CL motif analysis based on DEWseq results of ENO1 eCLIP (n=5). Logo representation of the top scoring motifs using DREME.

**Fig. S2.**
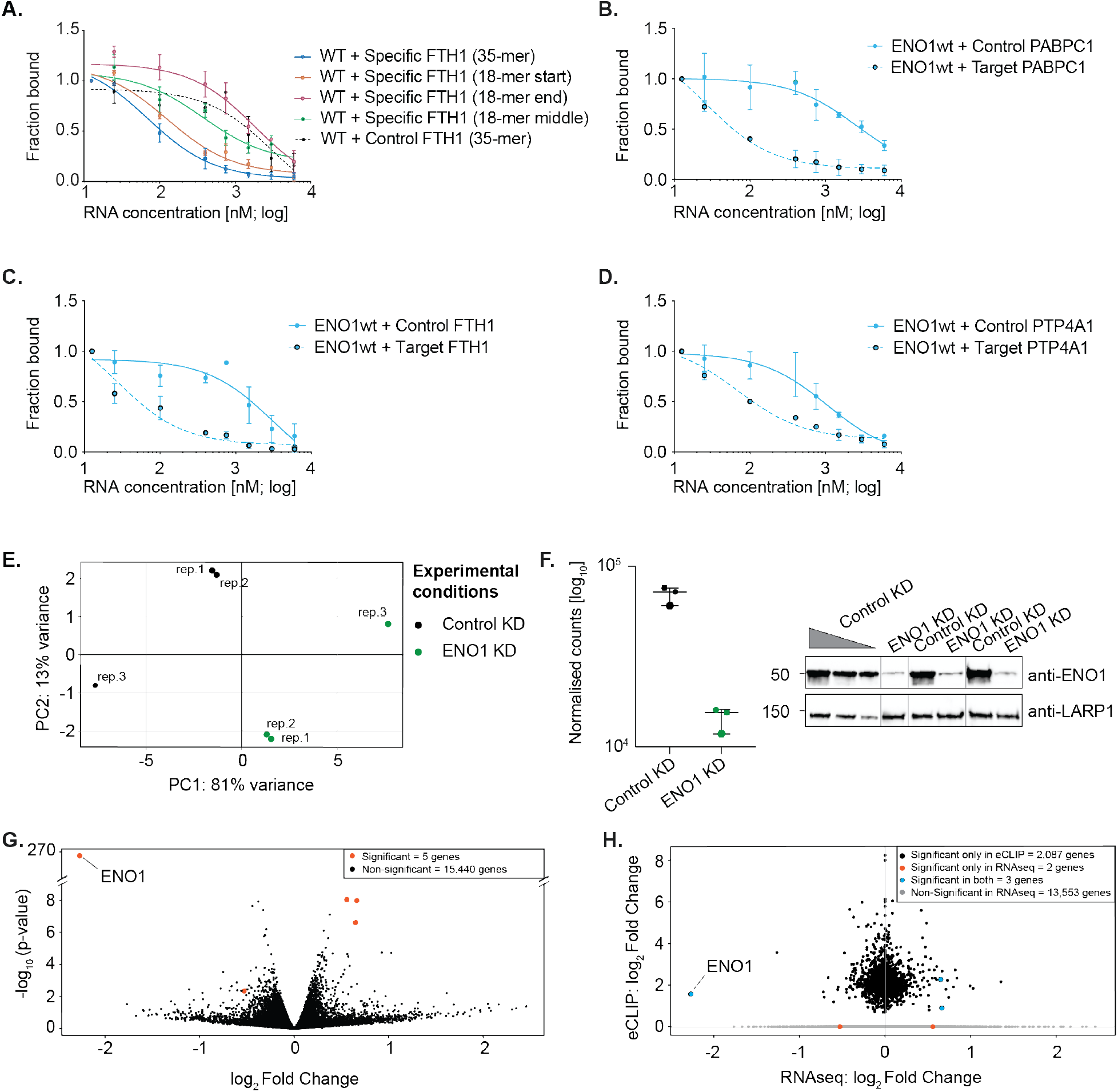
Specificity of ENO1 RNA-binding in vitro and transcriptome stability in HeLa cells after ENO1 knock-down. A.–D. In vitro analyses of ENO1 RNA binding specificity by competitive EMSAs. A. Quantification of competition EMSA experiments using FTH1 target RNA as a probe and different unlabeled 18-mer FTH1 RNA competitors (n=3). B. Quantification of competition EMSA experiments using PABPC1 target RNA as a probe and unlabeled PABPC1 control or target RNA competing for the binding to ENO1 (n=3). C. Quantification of competition EMSA experiments using FTH1 target RNA as a probe and unlabeled control or target FTH1 RNA as competitors (n=3). D. Quantification of competition EMSA experiments using PTP4A1 target RNA as a probe and unlabeled control or target (n=3). E.–H. Analysis of ENO1 knock-down (RNAi) effects on the HeLa cells transcriptome. E. PCA plot of biological replicates for control and ENO1 knock-down (KD) using RNAseq (n=3). F. Left: Normalized counts of RNA mapping to ENO1 mRNA after treatment with control or ENO1 siRNA. Right: Immunoblot of ENO1 and LARP1 for control and ENO1 KD. A dilution series (100, 50 and 25%) was loaded to assess the knock-down efficiency. G. Differentially expressed mRNAs upon ENO1 KD using RNAseq (p-value <0.1, log2 FC >0.5). H. Overlap of differentially expressed genes upon ENO1 KD and RNA-binding targets of ENO1 as detected in eCLIP (IHW-adjusted p-value <0.1, log2 FC >0.5; see Fig. 1C).

**Fig. S3.**
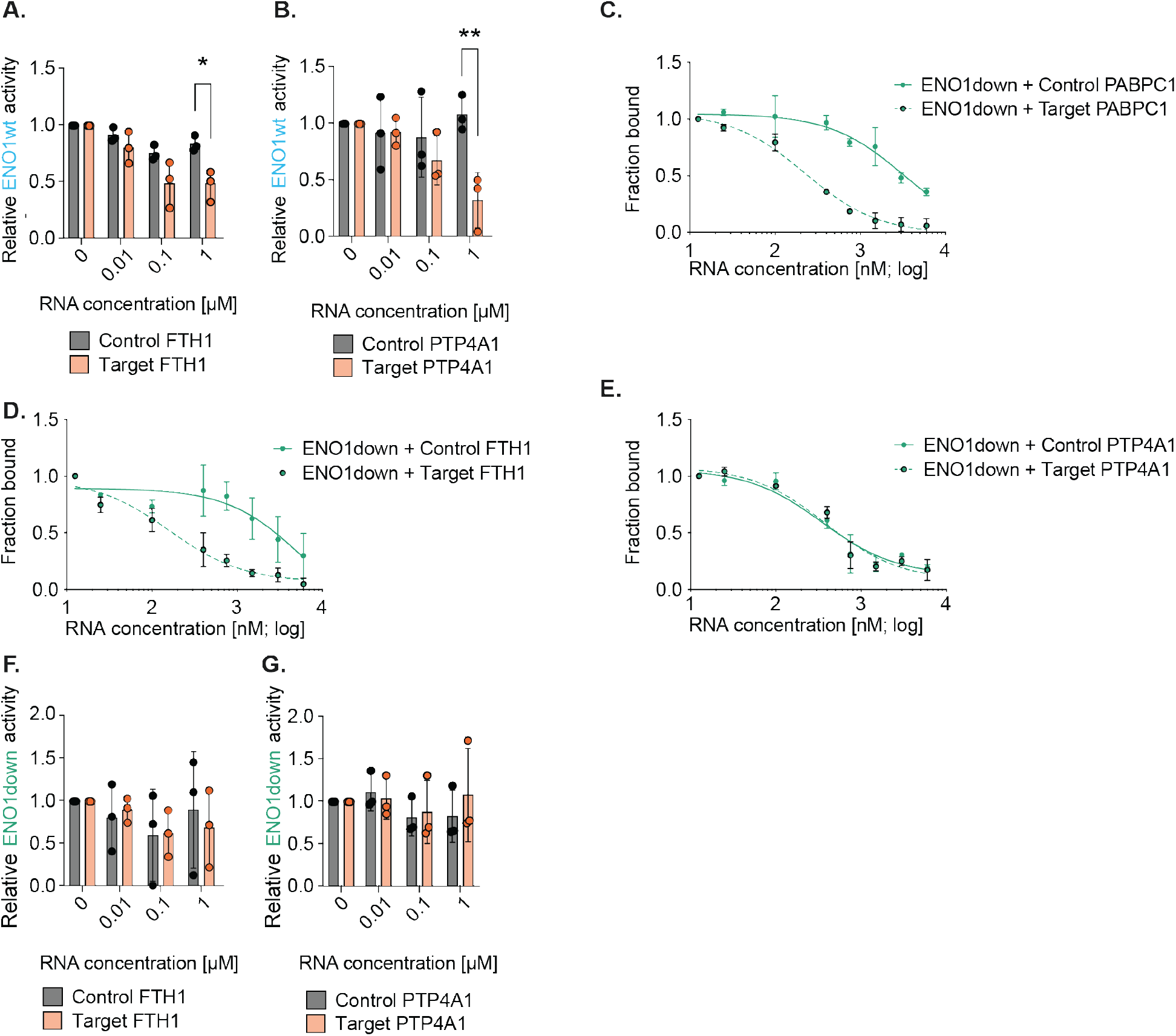
Riboregulation of ENO1 enzymatic activity in vitro. A. Measurement of recombinant ENO1wt activity with increasing concentrations of control and target FTH1 RNA (n=3). The standard deviation is given and the statistically significant differences were detected using two-way ANOVA and Sidak-correction for multiple comparison testing. B. Experimental setup as in A. for control and target PTP4A1 RNA. C. Quantification of competition EMSA experiments using PABPC1 target RNA as a probe and unlabeled PABPC1 control or target RNA competing for the binding to ENO1 (n=3). D. Quantification of competition EMSA experiments using FTH1 target RNA as a probe and unlabeled control or target FTH1 RNA as competitors (n=3). E. Quantification of competition EMSA experiments using PTP4A1 target RNA as a probe and unlabeled control or target (n=3) F. Measurement of recombinant ENO1down enzymatic activity with increasing concentrations of control and target FTH1 RNA (n=3). The standard deviation is given and the statistically significant differences were detected using two-way ANOVA and Sidak-correction for multiple comparison testing. G. Experimental setup as in F. for control and target PTP4A1 RNA.

**Fig. S4.**
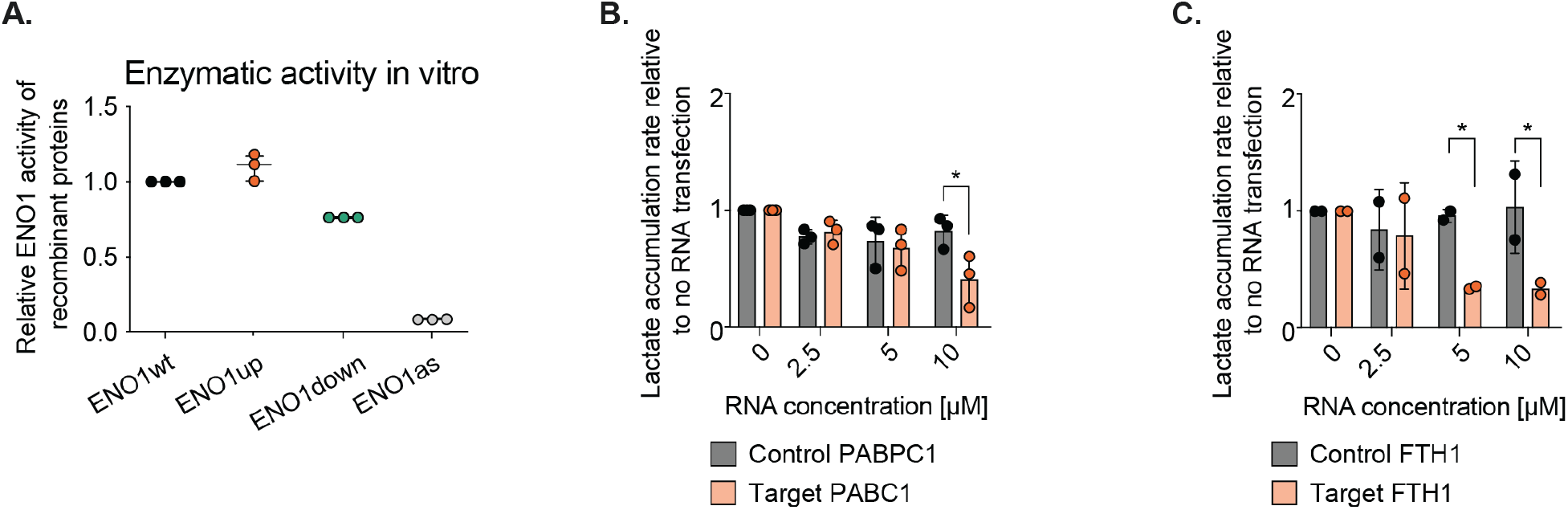
Riboregulation of ENO1 in HeLa cells. A. Relative enzymatic activity of ENO1wt, ENO1up, ENO1down and ENO1as determined with recombinant proteins in vitro (n=3). B. Increasing concentrations of target and control PABPC1 were nucleofected into HeLa cells. Upon nucleofection of the control or target RNAs, the lactate accumulation in the medium was measured after 30, 60 and 90 minutes and used to estimate the accumulation rate (n=3). The standard deviation is given and the statistically significant differences were detected using two-way ANOVA and Sidak-corrected for multiple comparison testing. C. Same experimental setup as in B. for FTH1 control and target RNA.

**Fig. S5.**
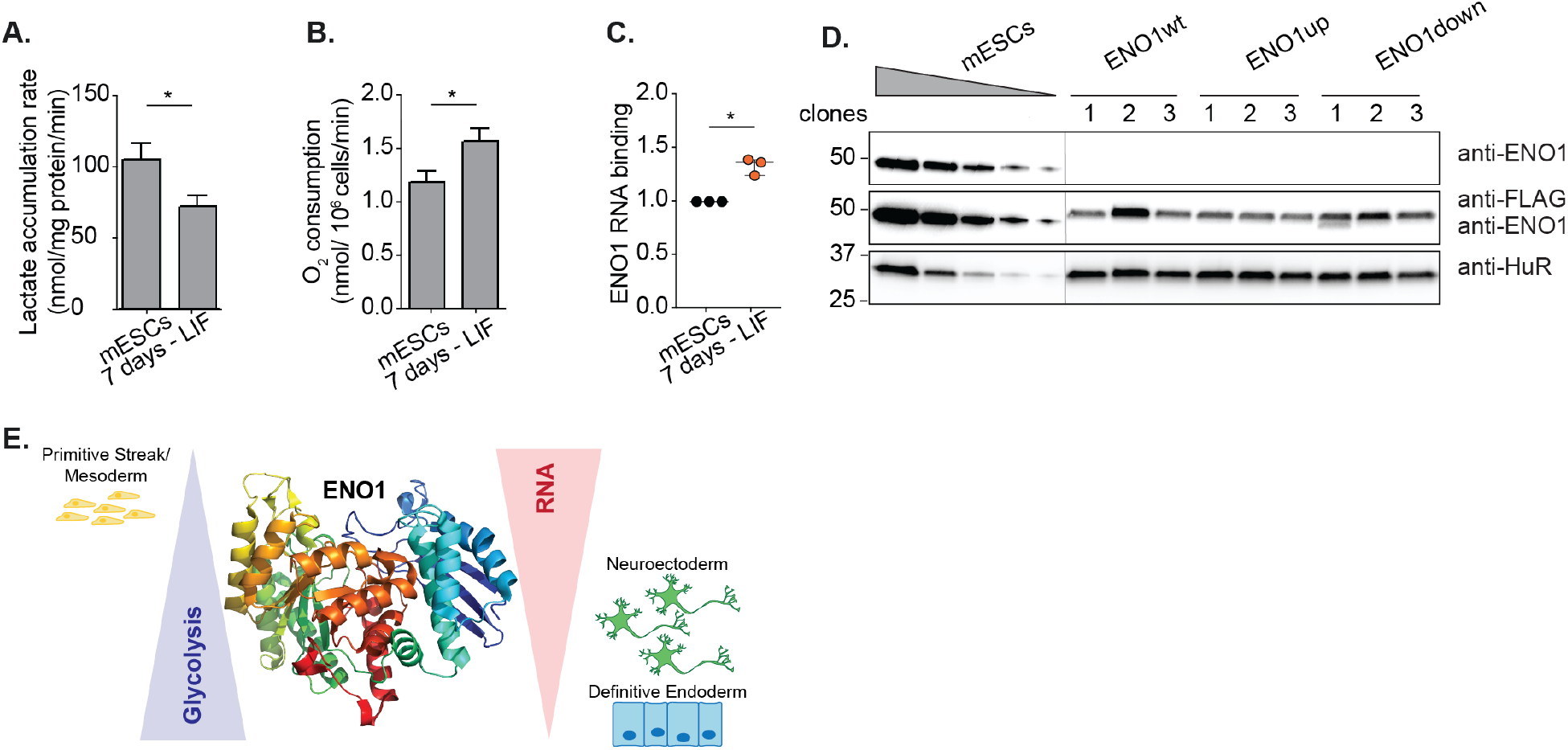
Analysis of ENO1 and mutant variants in mESCs. A. Lactate accumulation rate of pluripotent mESCs after LIF withdrawal for a period of seven days (n=3). The standard deviation is given and the two-tailed Student’s *t*-test is used to detect statistically significant differences. B. Oxygen consumption rate of cells was measured with the Oxytherm System (n=3). The standard deviation is given and the two-tailed Student’s t-test is used to detect statistically significant differences. C. Quantification of replicates of ENO1 PNK assays after UV-crosslinking for pluripotent mESCs and mouse cells after 7 days of LIF withdrawal (n=3). The standard deviation is given and the two-tailed Student’s *t*-test is used to detect statistically significant differences. D. Immunoblot of ENO1, FLAG and HuR (loading control) for three clones for ENO1 knockout and constitutive transgenic expression of Flag-HA-tagged ENO1wt, ENO1up and ENO1down from the Rosa26 locus. E. RNA riboregulates the enzymatic activity of ENO1, thereby inhibiting glycolysis and affecting mESC differentiation.

**Table S1.**
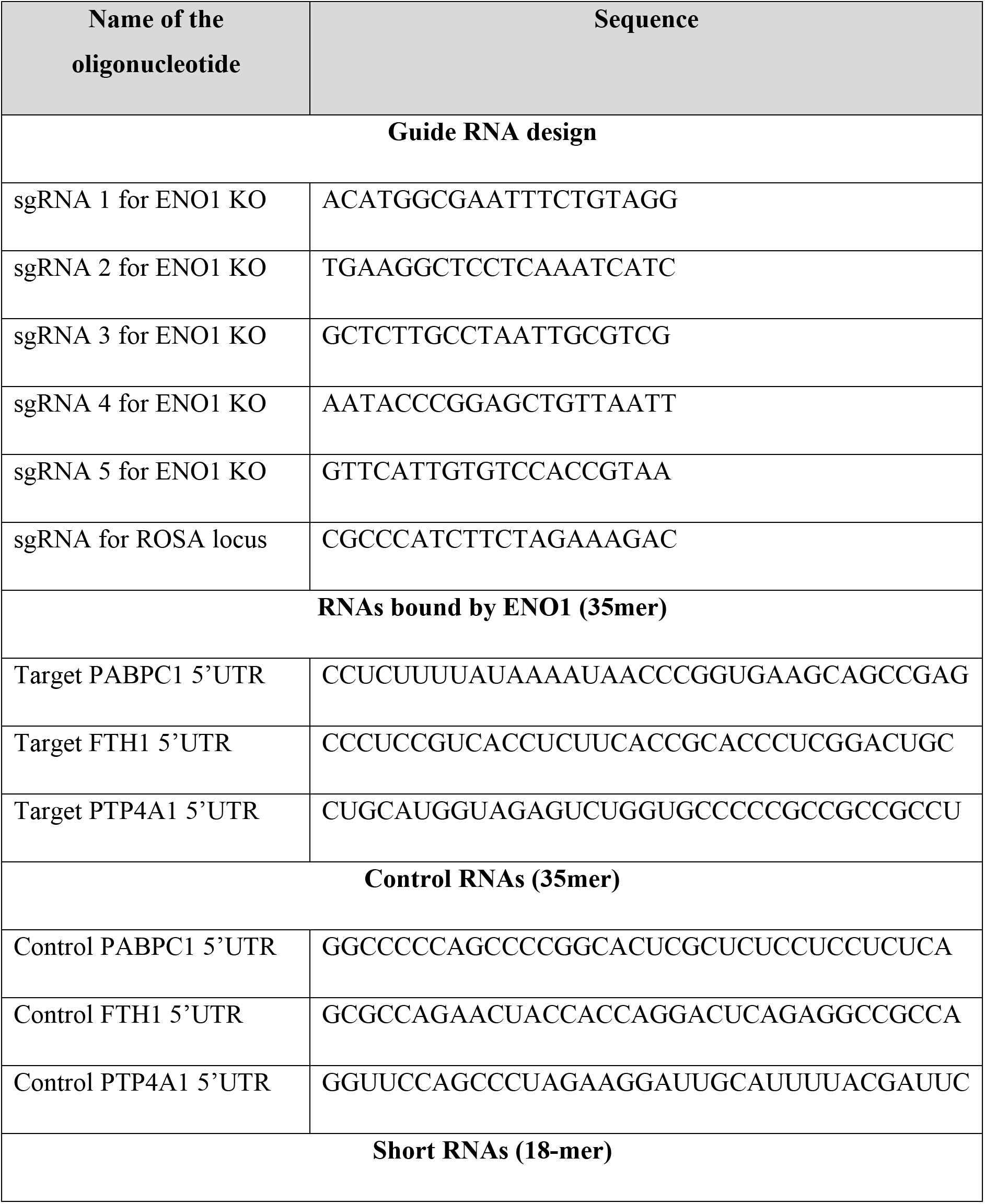
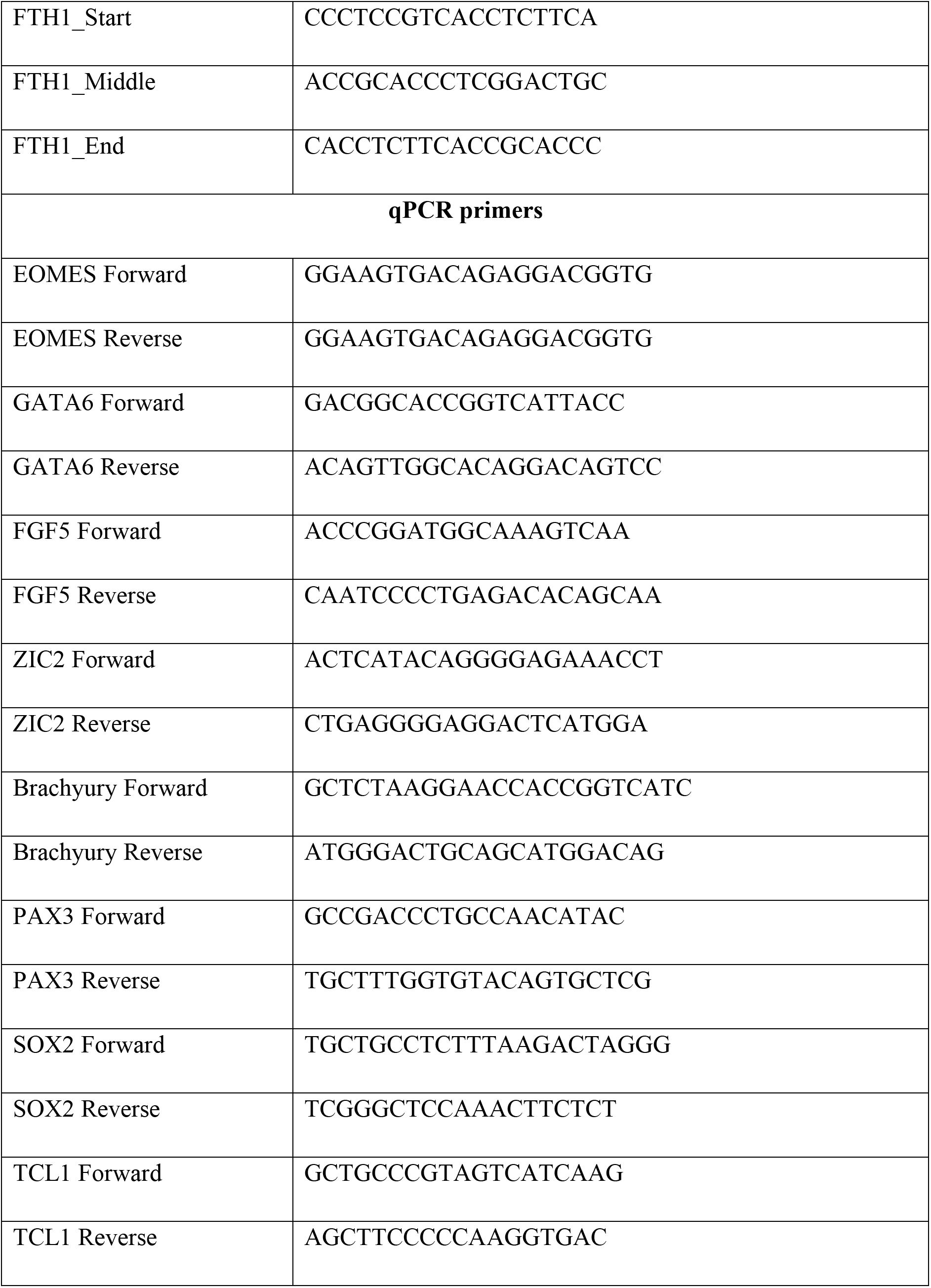
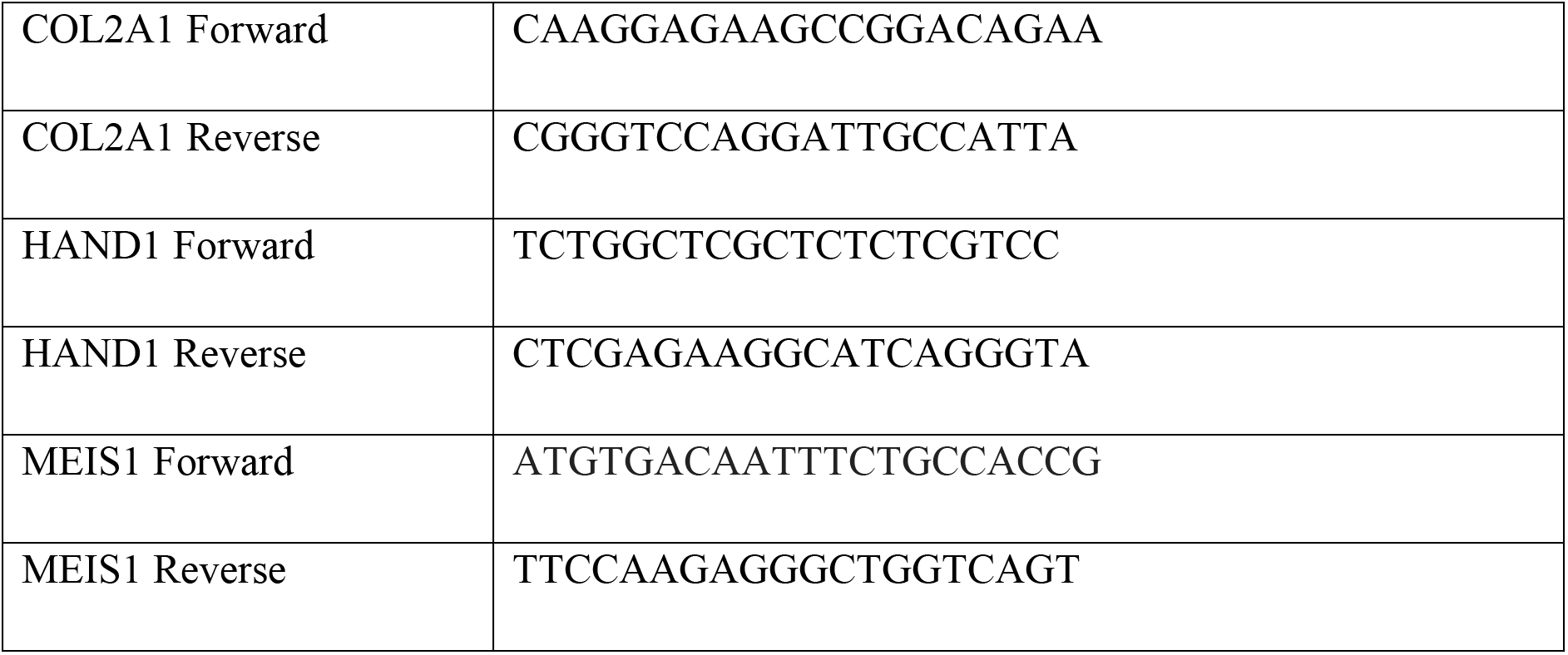
sgRNAs, qPCR primers and RNA sequences used in this manuscript

